# Tissue-resident memory T cells are diminished in the Kaposi sarcoma tumor microenvironment of people living with HIV

**DOI:** 10.64898/2026.10.10.758190

**Authors:** Anna Lagodka, Florian Winkler, Kveta Brazdilova, Robert Loewe, Katharina Grabmeier-Pfistershammer, Liv Eidsmo, Simona Saluzzo, Johanna Strobl

## Abstract

Kaposi sarcoma (KS) is a rare neoplasm and an AIDS-defining diagnosis in people living with HIV (PLWH). Although and chronic infection with Kaposi sarcoma–associated herpesvirus (KSHV) contributes to KS progression, the local immune mechanisms that are lost during tumor progression remain incompletely understood.

Here, we systematically characterized global patterns in the KS tumor microenvironment (TME) in both PLWH and HIV-negative individuals. Spatial transcriptomics revealed distinct niches comprising of tumor cells, immune cells, and surrounding skin cells. Tumor regions expressed a characteristic endothelial-fibroblastic signature attributable to KS tumor cells (CD34, LYVE1, SCN9A), TME regions expressed marker genes for T cells, B cells and cancer-associated fibroblasts. Analysis of putative cell-cell interactions demonstrated that tumor cells preferentially interact with cells of the TME, including T cells, indicating an important role as hub of cellular communication in KS.

Immunofluorescence imaging confirmed increased numbers of T cells, specifically tissue-resident memory T cells (T_RM_), in KS samples compared to healthy skin, accompanied by a pronounced upregulation of PD-1 expression that was predominantly localized to the TME. Notably, we identified HIV-associated alterations in the KS immune microenvironment: PLWH exhibited loss of CXCR3⁺ and IFN-γ–producing CD8⁺ T_RM_ cells in the KS TME compared to HIV-negative patients, a finding consistent with impaired antitumor immunity. Together, our findings reveal HIV-associated disruption of functional anti-tumor immunity in the KS TME of PLWH. Increased PD-1 expression within T cells of the TME highlights potential therapeutic use of immune checkpoint inhibitors in KS.

**What is already known on this topic:** While systemic immunosuppression and viral replication are known drivers of KS progression, the local, microanatomical immune mechanisms governing tumor control remain under-investigated. Specifically, how HIV infection functionally alters antitumor immunity within the local tumor TME and why PLWH frequently exhibit impaired antitumor responses compared to HIV-negative individuals, has not been investigated.

**What this study adds:** This study provides a spatially resolved cellular and transcriptomic map of the KS microenvironment across both PLWH and HIV-negative cohorts, indicating that T cells in the KS TME serve as a primary hub of cellular communication. Crucially, we discovered an HIV-specific loss of functional, CXCR3+CD8+ T_RM_ cells in the TME. Lastly, we identify activity of co-inhibitory receptor pathways and high PD-1 expression in KS-associated T cells.

**How this study might affect research, practice or policy:** These findings advance tumor immunology by defining functional impairment and local depletion of T_RM_ as a hallmark of HIV-associated malignancy. Furthermore, the pronounced upregulation of PD-1 within the TME provides a strong biological rationale for the use of immune checkpoint inhibitors in patients with KS.

**Graphical Abstract:** KS lesions are populated by cytokine-producing CXCR3+ PD-1+ T cells. TME in HIV-associated KS is characterized by loss of IFN-g-producing T_RM_ leading to impaired functional tumor-immunity. (Created in BioRender. Strobl, J. (2026) https://BioRender.com/drwi6gk.)

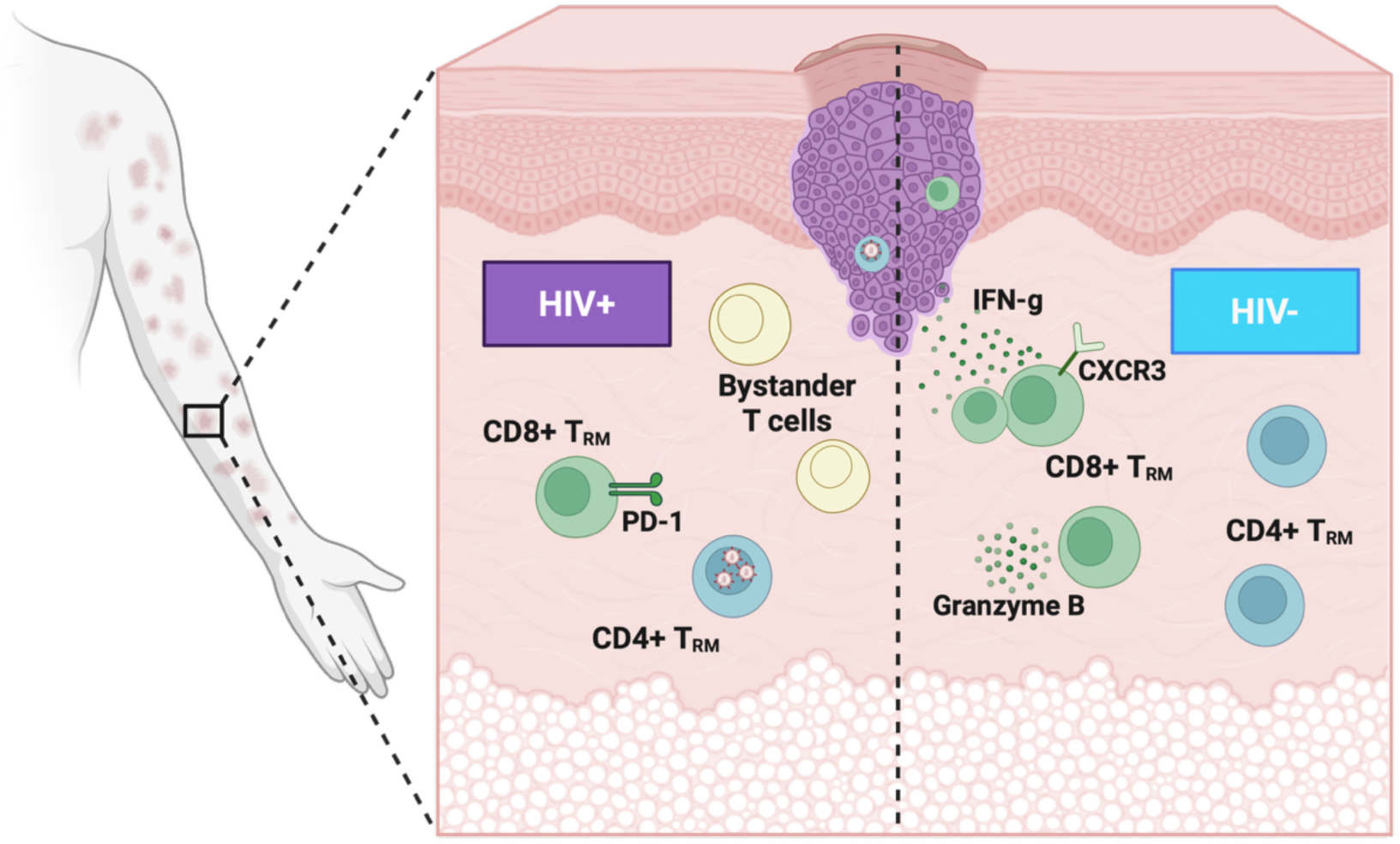

## Introduction

Kaposi sarcoma (KS), which occurs in people with human immunodeficiency virus (HIV) infection, is both an AIDS defining disease and the most common neoplasm in people living with HIV (PLWH) (1, 2). In addition to HIV-infected individuals, KS may also affect immunocompromised individuals and prevalence is increased in KS associated herpesvirus (KSHV) endemic areas (3). The causative agent is discussed as Kaposi sarcoma-associated herpesvirus (KSHV), also known as Human herpesvirus 8 (4).

Histologically, cutaneous KS lesions are characterized by the proliferation of KSHV-infected spindle cells of endothelial origin, accompanied by abnormal vascular structures, inflammatory infiltrates, and fibrosis (5). Clinically, lesions present as violaceous macules, papules and nodules, or progress with edema and ulcerations, or even to disseminated, multi-organ disease, particularly in immunocompromised individuals (5, 6).

Depending on etiology and the affected patient cohort, KS is classified into several subtypes: The classic form predominantly arises in elderly men of Mediterranean or Eastern European origin and is generally characterized by a slow and indolent course; endemic KS is prevalent in sub-Saharan Africa and often involves lymph nodes and internal organs; iatrogenic KS develops in immunosuppressed patients, most commonly following solid organ transplantation; epidemic (HIV-associated) KS is an AIDS-defining disease that often presents with extensive skin, mucosal, lymph node, and organ manifestations (6). A fifth, newly described subtype is non-epidemic KS in HIV-negative men who have sex with men (MSM). This variant resembles the classic form but typically affects younger individuals and is generally associated with a more benign course (7).

As in many other neoplasms, decreased immunosurveillance and chronic virus infection seem to play an important role in KS progression. KSHV is known to promote a pro-inflammatory tumor microenvironment (TME) that encompasses the recruitment of myeloid and lymphoid immune cells (8). The TME is rich in natural killer cells (NK) and CD8⁺ T cells as well as high levels of CXCL12 and interferon-gamma (IFN-γ) with distinct differences in immune cell composition and cytotoxic signatures observed between HIV-positive and HIV-negative patients Whole-transcriptome analyses of FFPE sections demonstrated an upregulation of NK-cell-mediated immunity and cytotoxic responses in HIV-negative KS, as contrasted with stronger signatures associated with cell proliferation and protein kinases in HIV-positive KS. However, the spatial organization of these distinct immune states within KS lesions remains poorly understood. (9). It has, however, been demonstrated that despite the presence of T cell– attracting chemokines, such as CXCL9 and CXCL12, CD8⁺ T cells are frequently excluded from areas with high densities of KSHV-infected cells, indicating a potential mechanism of immune evasion (10). KSHV further modulates the immune response by inhibiting CD4⁺ T cell proliferation and skewing immunity toward a Th2-dominant response (11, 12).

Skin-resident T cells, especially CD8⁺ T_RM_, have been shown to be important mediators of local tumor control in skin cancer and other solid malignancies. T_RM_ cells are defined by the expression of CD69, CD103, and/or CD49a and reside permanently within tissues, where they mediate rapid, site-specific immune responses (13, 14). Saluzzo et al. recently showed that delayed antiretroviral therapy (ART) initiation results in irreversible depletion of T_RM_ cells in skin and mucosa, potentially resulting in increased anal cancer risk. This was accompanied by a significantly reduced expression of CXCR3 on cutaneous CD4⁺ and CD8⁺ T_RM_ cells, suggesting not only a quantitative loss of T_RM_ cells but also a qualitative change in local Th1/cytotoxic immunity and T-cell recruitment to inflamed tissues (15). However, KS also develops in individuals with normal T cell counts, while the contribution of tissue-resident memory T cells has not been addressed. It remains unknown if and how lack of local cellular immunity leads to KS tumor progression and there is active research on how this can be targeted therapeutically (16). The present study therefore assessed the KS TME in HIV-positive and HIV-negative individuals with a focus on skin T_RM_ cells using immunophenotyping, imaging and spatial RNA-sequencing.

## Methods

### Patient recruitment and sampling

Recruitment included pre-treatment skin biopsies from individuals with cutaneous Kaposi sarcoma (KS) in two groups: PLWH, HIV-positive patients with a low nadir (n=6), and HIV-negative patients with cutaneous KS (n=5-6). All KS diagnoses were confirmed by expert histopathologists at the Medical University of Vienna, Department of Dermatology. Patient characteristics are detailed in Supplementary Table 1.

### Tissue processing and imaging

Biopsies were processed as either cryosections or formalin-fixed paraffin-embedded (FFPE) sections. Tissue was sectioned at 7 μm thickness. Cryosections were mounted on glass slides, fixed in acetone at 4°C for 10 minutes, and stored at −20°C until staining. Prior to staining, FFPE sections were deparaffinized (60°C), followed by Neo-Clear and ethanol treatments, and heat-induced antigen retrieval. Frozen slides were equilibrated to room temperature and washed in phosphate-buffered saline (PBS) for 10 minutes. For multicolor immunofluorescence staining, sections were blocked with 3% mouse serum in 2% BSA/PBS for 20 minutes to minimize nonspecific binding. Directly labeled monoclonal antibodies targeting relevant markers (e.g., CD4, CD8,) were applied in 2% BSA/PBS for 2 hours at room temperature. After washing in PBS, nuclear counterstaining was performed with DAPI in 2% BSA for 4 minutes. Sections were washed again and mounted with antifade medium. Image acquisition was performed using a Z1 Axio Observer microscope (Zeiss) with an LD Plan-Neofluar 20×/0.4 objective. Images were captured under standardized exposure settings and analyzed using TissueQuest software (TissueGnostics) for quantitative assessment of marker expression and immune cell localization.

### Statistical analyses

Statistical analysis included a minimum of six patients per group, with all experiments performed in duplicate. Quantitative data were compared using one-way ANOVA with Tukey’s post hoc test for multiple comparisons. Statistical significance was defined as P < 0.05. Data analysis and visualization were performed using GraphPad Prism version 8.

### Sample preparation and downstream analysis for spatial transcriptomic profiling

To examine the spatial relationships among cell types involved in cell–cell communication, spatial transcriptomics was performed using the Visium platform (10x Genomics), as previously described (17). KS samples were embedded in Tissue-Tek® optimum cutting temperature (O.C.T.™) compound (Sakura Finetek, Torrance, CA, USA) and snap-frozen in liquid nitrogen. Sections (10 µm) were cut, placed onto a Visium Spatial Gene Expression Slide, and processed following the manufacturer’s protocol, including methanol fixation, hematoxylin–eosin staining, and imaging. Subsequent permeabilization and library preparation were carried out according to the Visium (10x Genomics) protocol and sequenced by the Biomedical Sequencing Facility at the CeMM Research Center for Molecular Medicine of the Austrian Academy of Sciences, using the Illumina NovaSeq 6000 platform. Raw sequencing data were processed using the Space Ranger pipeline (v1.2.0, 10x Genomics). Aligned data processing, cell type annotation, and communication analysis were performed using the Python toolkit SCANPY (18) and R package Cellchat (version 1.6.1) (19).

### Ethics Statement

All authors have declared that no conflict of interest exists.

## Author Contributions Statement (CRediT-compliant)

A.L.: methodology, investigation, data analysis, writing (original draft); F.W.: methodology, investigation, data analysis, writing (original draft); R.L.: investigation, supervision. K.B.: investigation, data analysis; K.G.P., L.E.: investigation, supervision. S.S.: methodology, supervision; J.S.: Conceptualization, funding acquisition, methodology, investigation, data analysis, writing (original draft); All authors: writing (review & editing).

## Declaration of Generative AI Use

During the preparation of this work, the author(s) used *OpenEvidence* to support literature exploration and to assist in summarizing scientific background information. After using this tool, the authors critically reviewed, verified, and edited all generated content as needed and take full responsibility for the accuracy and integrity of the published article.

## Results

To define distinct tissue niches and identify the immune and stromal components that shape microenvironmental niches in KS, we performed spatial transcriptomic profiling of KS lesional biopsies from two HIV+ individuals. We detected distinct tissue niches through Leiden clustering based on established marker genes (Figure 1A), allowing the identification of tumor cell-rich regions, and regions dominated by tumor-microenvironment (TME)–associated cells, activated endothelial cells, cancer-associated fibroblasts, plasma cells and keratinocytes. The corresponding violin plots demonstrated that the KS markers *CD34*, *LYVE1* and *SCN9A* (20) were predominantly expressed in tumor cell areas (Figure 1B). *CD34* additionally showed strong expression in endothelial cell regions and, to a lesser extent, in activated and cancer-associated fibroblasts, reflecting the strong endothelial–fibroblastic signature typical for KS lesions (20). Using a combinatory KS tumor gene signature (co-expression of *LYVE1, SCN9A, CD34, PECAM1, PDPN, FLT4*), we mapped KS tumor regions to histology images (Supplementary Fig. 1A, B).

**Figure 1.**
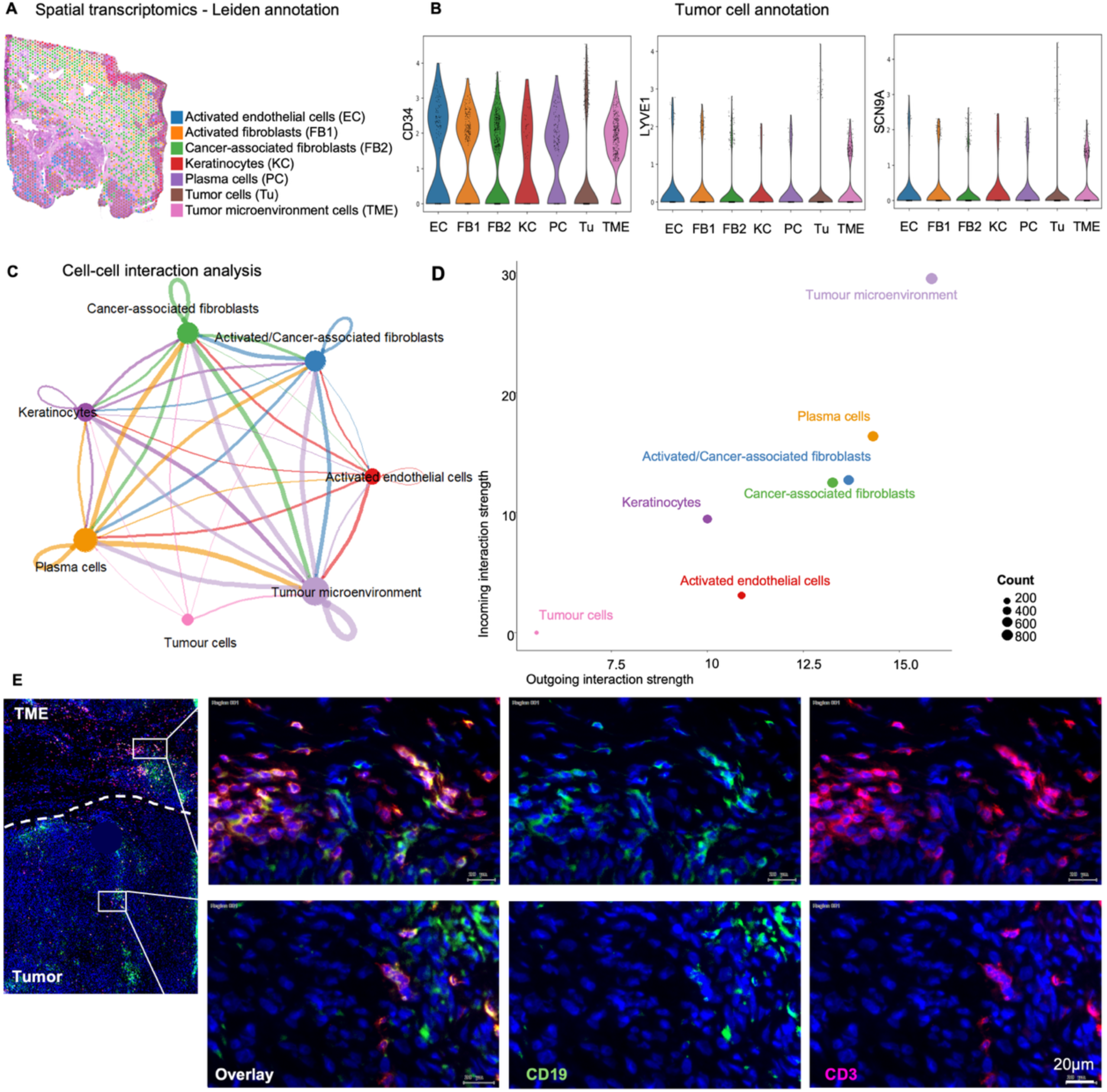
**Spatial transcriptomic mapping of immune cell interactions in KS**. (A) Broad cell type annotation by Leiden clustering using known marker genes from spatial transcriptomics (Visium) of a KS sample. (B) Violin plots showing gene expression of CD34 (left), LYVE1 (middle) and SCN9A (right) across cell types from (A). (C) Circle plot displaying the aggregated cell–cell communication network, with edge with representing the strength of interactions between cell types and node size representing the total number of outgoing interactions. (D) Scatter plot of incoming (y-axis) and outgoing (x-axis) communication probabilities for each cell type from (A); dot size represents the number of inferred links. (E) Representative immunofluorescence images of CD3⁺ T cells (red) as well as CD19⁺ B cells (green) with nuclear counterstaining using DAPI (blue) in KS tissue sections. Scale bar = 20 μm.

We next investigated the communication network between these diverse cellular compartments using the R package CellChat. Unsupervised analysis of overall intercellular communication revealed a strong outgoing and incoming interaction probability for the tumor microenvironment (TME, including *CD3E-* and *ITGAE-*expressing T cells, Supplementary Fig. 1C), suggesting that the TME represents the dominant site of cellular communication in KS (Figure 1C-D). Plasma cell as well as activated and cancer-associated fibroblast-rich regions also exhibited strong outgoing interaction probabilities. In contrast, areas populated by keratinocytes and endothelial cells displayed lower outgoing and incoming interaction probabilities, while tumor cells showed the lowest overall interaction probability (Figure 1C). Immunofluorescence imaging demonstrated infiltration of CD3+ T cells and sparse CD19+ B cell infiltrates in the TME surrounding tumor regions (Figure 1E). Quantification of B cells did not reveal significant differences compared to healthy skin (Supplementary Figure 1D). However, we detected significant T cell enrichment in the KS TME (Figure 2A, B).

**Figure 2.**
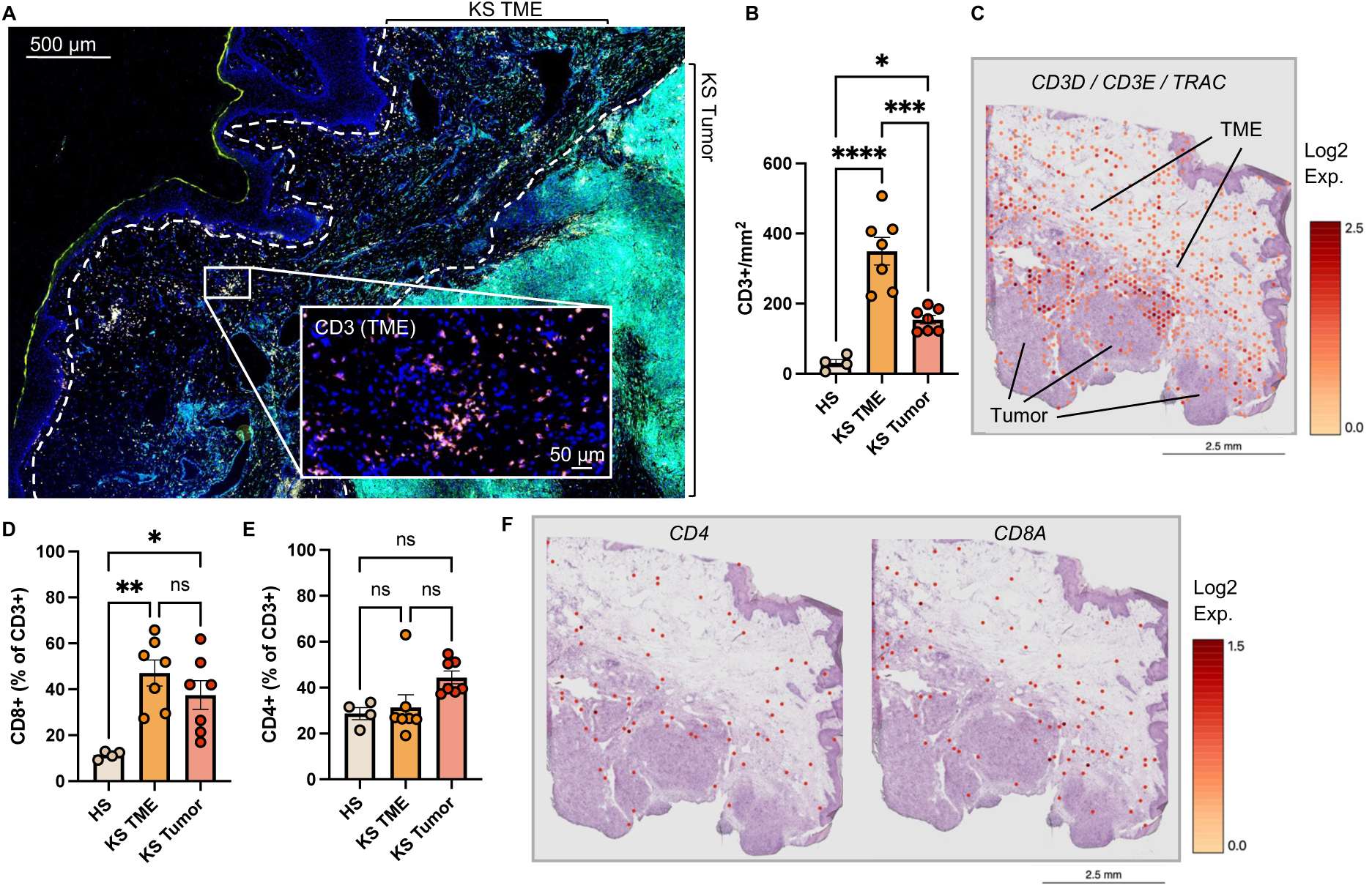
Predominance of cytotoxic T cells in the KS TME. (A) Representative immunofluorescence image CD3⁺ T cells (magenta) with nuclear counterstaining using DAPI (blue) in KS tissue sections. Tumor-autofluorescence seen as cyan. (B) Total number of T cells/mm^2^ (CD3⁺) in healthy skin (HS, n=4), the KS tumor microenvironment (TME, n=7), and intratumoral areas (Tumor, n=7) of KS lesions. (C) Representative spatial gene expression mapping of combined *CD3D, CD3E* and *TRAC* expression (log2) on an H&E-stained KS tumor sample. (D-E) Quantification of (D) cytotoxic T cells (CD3⁺CD8⁺) and (E) helper T cells (CD3⁺CD4⁺) in HS, KS TME, and tumor compartments. Data shown as percentage of CD3+ cells. (F) Representative CD4 and CD8A gene expression (log2) in the sample from (C). Data are presented as individual values and mean ± SEM. Statistical analysis was performed using one-way ANOVA. *P < 0.05; **P < 0.01; ***P < 0.001, ****P<0.0001.

Together, these spatially resolved datasets suggest that the TME is the main site of coordinated immune and stromal interactions, laying the basis for subsequent analyses focusing on T cell patterns in the KS TME.

T cells was a prominent immune cell type in the KS TME, and we next sought to validate our initial analyses using tissue immunofluorescence of a larger KS tumor cohort. Image-based quantification revealed significantly increased infiltration of CD3⁺ T cells in KS tissue compared to healthy skin (Figure 2A, B). In spatial transcriptomics data, T cell-associated genes *CD3D*, *CD3E* and *TRAC* were enriched in peri-tumoral regions (Figure 2C). T cell enrichment was most prominent in the peritumoral areas (TME), with only modest increase of tumor-infiltrating CD3⁺ T cell numbers, indicating that T cells do not efficiently infiltrate the tumor mass. When assessing specific T cell subsets, we observed a marked increase of CD8⁺ T cells in KS samples, especially in the TME (Figure 2D). In contrast, CD4⁺ T cells remained relatively stable across disease states and tumor compartments (Figure 2E). Spatial transcriptome data confirmed these findings by revealing specific regions with high *CD8A* and *CD4* gene expression, particularly within peritumoral zones (Figure 2F, Supplementary Figure 2A).

Tissue-residing T cells play a pivotal role in controlling solid tumors. To assess the presence, phenotype, and distribution of T_RM_ in KS, we quantified CD69^+^ and CD103^+^ T cells in different tissue compartments. Both T_RM_ markers were significantly increased in KS lesions compared to healthy skin (Figure 3A) and present on transcriptional level (*CD69*, *ITGAE*, Figure 3B). Notably, CD8^+^ T_RM_ cells were particularly enriched in the TME, while no significant differences in CD4^+^ T_RM_ cell frequencies were observed between HS, TME, and tumor tissue (Figure 3C, D). In addition, increased numbers of tumor-infiltrating CXCR3^+^ T_RM_ cells were observed in KS tissue, with a significant accumulation of CXCR3^+^CD8^+^ T cells in the tumor microenvironment (Figure 3E, F). Spatial transcriptomic analysis confirmed these findings, demonstrating expression of *CXCR3* and associated recruitment genes *CXCL9* and *CXCL10* not only in the tumor and TME, but also in the surrounding non-lesional skin, indicating a broader immune imprint (Figure 3G, Supplementary Figure 2A).

**Figure 3.**
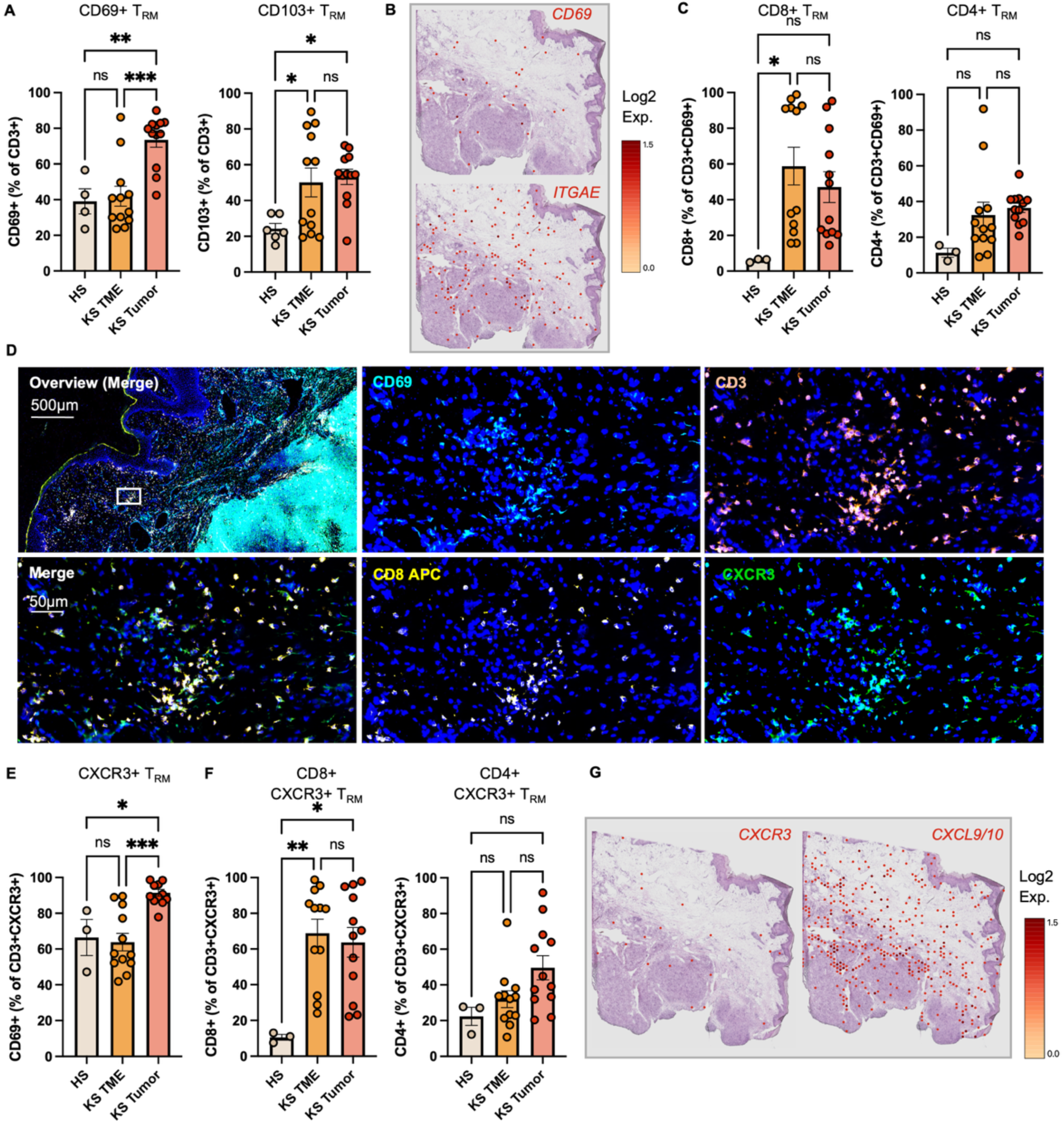
Enrichment of CXCR3⁺ TRM cells in KS lesions. (A) Quantification of TRM (CD3⁺CD69⁺ and CD3⁺CD103^+^) in HS (n=4), KS (n=11) TME and intratumoral areas (tumor) from IF data. Data shown as percentage of CD3⁺ cells. (B) Representative spatial gene expression plot of *CD69* and *ITGAE* log2-expression in a KS tumor sample. (C) Quantification of cytotoxic and helper TRM (CD8⁺ and CD4⁺) in samples from (A). Data shown as percentage of TRM (CD3⁺CD69⁺). (D) Representative IF image of CD69, CD3, CD8, and CXCR3, and merge with nuclear counterstaining using DAPI (blue), in the TME of KS tissue sections. Scale bar = 500 µm (overview) and 50 μm (detail). (E-F) Quantification of CXCR3⁺ TRM (E, CD3⁺CXCR3⁺CD69⁺), CXCR3⁺ cytotoxic T cells (F, CD3⁺CXCR3⁺CD8⁺) and CXCR3⁺ helper T cells (F, CD3⁺CXCR3⁺CD4⁺) in samples from (A). Data shown as percentage of the indicated cell type. (G) Representative spatial transcriptomic profiling illustrating distribution of *CXCR3* and combined *CXCL9/CXCL10* gene expression densities across compartments. In (A), (C) and (E-F) data are presented as individual values and mean ± SEM. Statistical analysis was performed using one-way ANOVA. *P < 0.05; **P < 0.01; ***P < 0.001.

Together, these results indicate a strong presence of cytotoxic CXCR3^+^ T_RM_ populations in the local immune landscape of KS, implicating them in anti-tumor immune responses.

The effector profile of T cells and T_RM_ in KS lesions was assessed next. Immunofluorescence-based cytokine detection and transcriptomic analyses revealed high frequencies of Granzyme B (GzmB)–positive T cells across KS tissue compartments (Figure 4A, B, Supplementary Figure 2A). GzmB-producing T_RM_ cells were also abundantly present, indicating cytotoxic potential within the resident immune compartment. IFN-γ–producing T cells were observed at lower levels overall, with increased expression was seen in the TME of KS samples (Figure 4C, D). Spatial transcriptomic mapping further supported widespread distribution of *GZMB*-expressing areas around the tumor. While only low numbers of *IFNG* transcripts were detectable, we observed a strong *STAT1* gene signature in KS peritumoral (TME) areas (Figure 4E, Supplementary Figure 2A).

**Figure 4.**
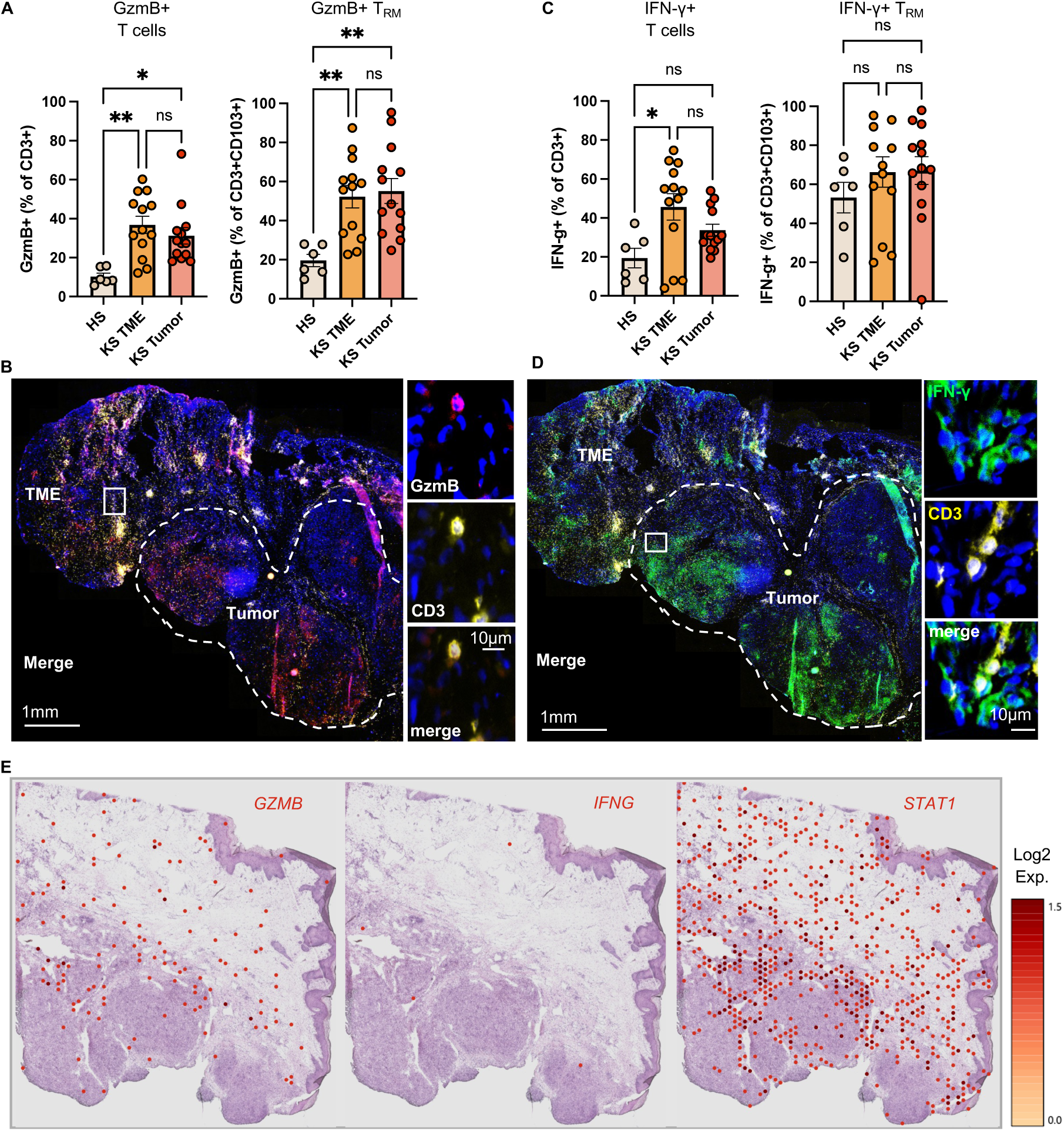
Abundance of cytokine-producing cytotoxic TRM cells in KS lesions. (A) IF-based quantification of GzmB-producing T cells (CD3⁺GzmB⁺) and GzmB-producing TRM (CD3⁺CD103⁺GzmB⁺) in HS (n=6), KS (n=11) TME, and tumor compartments. Data shown as percentage of T cells (CD3⁺) and TRM (CD3⁺CD103⁺). (B) Representative IF images showing GzmB⁺ cells (red), CD3⁺ T cells (yellow) and merge with nuclear counterstaining using DAPI (blue), in KS tissue sections. Scale bar = 1mm (overview image) and 10μm (detail). (C) Quantification of IFN-γ producing T cells (CD3⁺IFN-γ⁺) and IFN-γ producing TRM (CD3⁺ CD103⁺IFN-γ⁺) in samples from (A). Data shown as percentage of T cells (CD3⁺) and TRM (CD3⁺CD103⁺). (D) Representative immunofluorescence images showing IFN-γ⁺ cells (green), CD3⁺ T cells (yellow) and merge with nuclear counterstaining using DAPI (blue), in KS tissue sections. Scale bar = 1mm (overview image) and 10μm (detail). (E) Spatial gene expression plots (log2) illustrating *GZMB*, *IFNG* and *STAT1* expression densities across compartments. In (A) and (C), data are presented as mean ± SEM. Statistical analysis was performed using one-way ANOVA. *P < 0.05; **P < 0.01; ***P < 0.001.

Together, these findings demonstrate that KS lesions are enriched with cytotoxic T_RM_ cells, and to a lesser extent IFN-γ–producing subsets, reflecting a functionally activated T-lymphocytic immune infiltrate that is mostly confined to tumor-surrounding areas.

Since HIV-infection is associated with persistent depletion of skin T_RM_ cells following delayed ART initiation (15), we hypothesized that HIV-positive individuals had an altered local immune immune landscape in KS and directly compared T cell infiltration patterns between HIV-positive patients with low nadir and HIV-negative patients.

HIV-positive individuals exhibited higher overall CD3⁺ T cell infiltration in the TME, but no significant increase in tumor-infiltrating lymphocytes was observed (Figure 5B). In contrast, HIV-negative patients showed significantly higher levels of tumor-infiltrating CD8⁺ T cells (Figure 5B-D). As expected, CD4⁺ T cells were decreased in the TME of HIV-positive patients, while no significant differences in the frequencies of tumor-infiltrating CD4⁺ T cells were observed between groups (Figure 5C). Notably, we also found that HIV-positive patients harbored significantly lower levels of both CD69+ and CD103+ T_RM_ in the TME (Figure 5E). CD8⁺ T_RM_ cells were drastically reduced in both the tumor and TME compartments of HIV-positive individuals (Figure 5F). Furthermore, the total number of CXCR3⁺ T cells was reduced in the HIV-positive cohort and a more refined analysis revealed that CXCR3⁺CD8⁺ T cells were significantly depleted in both the TME and tumor of PLWH (Figure 5G-H).

**Figure 5.**
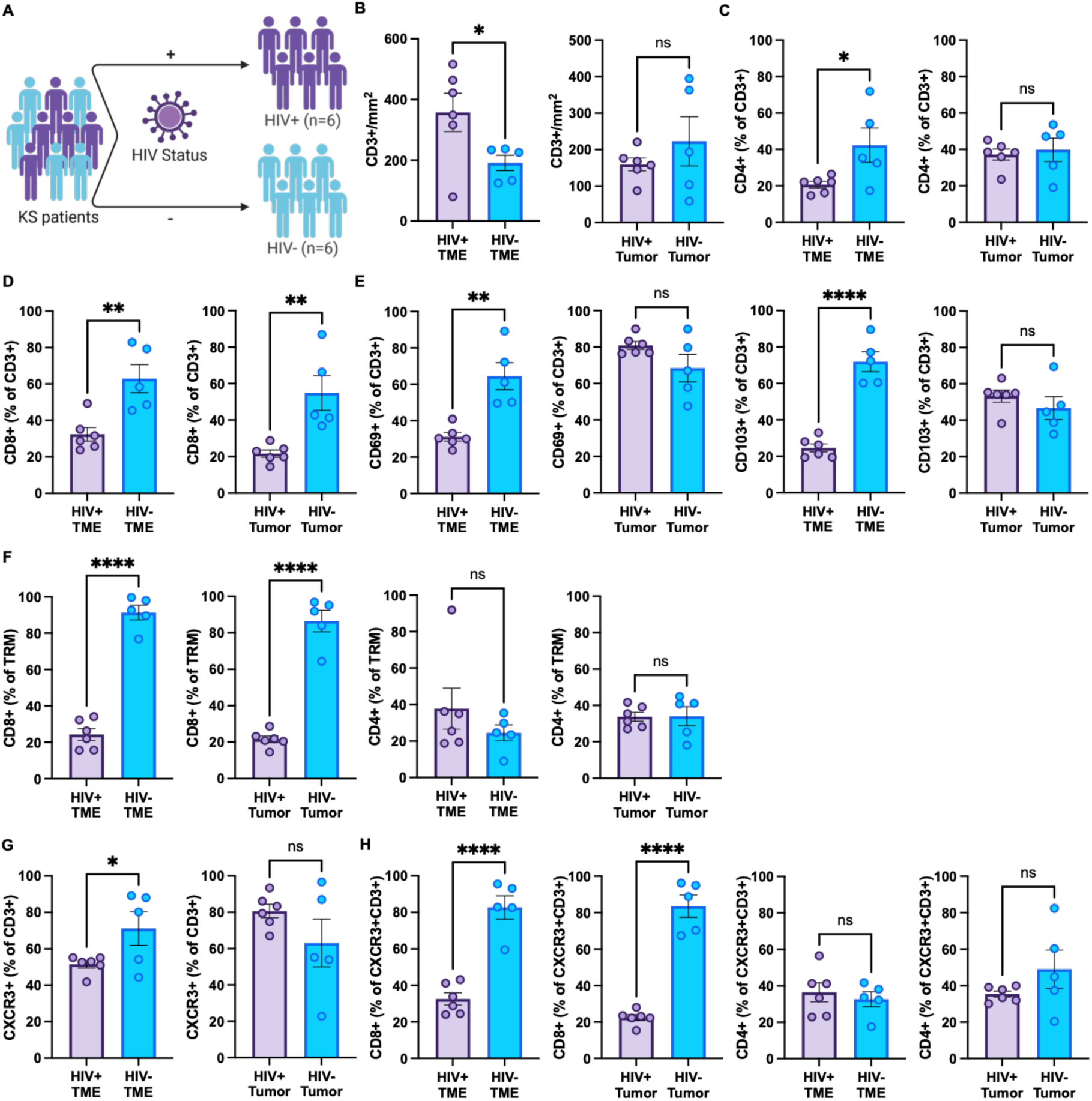
**Distinct immune cell infiltration patterns in KS tissue of HIV+ and HIV-patients**. (A) Graphical representation of patient cohorts. (B) Total number of T cells (CD3+) in the TME and tumor compartments of HIV+ (n=6) and HIV-patients (n=5) with KS. (C-D) Quantification of helper T cells (CD3⁺CD4⁺, C), and cytotoxic T cells (CD3⁺CD8⁺, D) in samples from (B). Data shown as percentage of T cells (CD3⁺). (E) Quantification of TRM (CD3⁺CD69⁺, CD3⁺CD103⁺) in in samples from (B). Data shown as percentage of T cells (CD3⁺). (F) Quantification of CD8⁺ and CD4⁺ subsets among CD3⁺CD69⁺ TRM in samples from (B). (G) Quantification of CD3⁺CXCR3⁺ T cells (CD3⁺CXCR3⁺) in samples from (B). Data shown as percentage of T cells (CD3⁺). (H) Quantification of CXCR3⁺CD8⁺ and CXCR3⁺CD4⁺ T cells in samples from (B). Data shown as percentage of CXCR3⁺ T cells (CD3⁺CXCR3⁺). In (B)-(H) data are presented as mean ± SEM. Statistical analysis was performed with unpaired Student’s t test. *P < 0.05; **P < 0.01; ****P < 0.0001.

Together, these data indicate that HIV-associated KS is characterized by depletion of CXCR3⁺CD8⁺ and CD8⁺ TRM populations, consistent with impaired tissue-resident immune surveillance in the local tumor microenvironment.

To further characterize functional T cell states, we analyzed effector molecule expression in T cells across PLWH and HIV-negative KS patients. GzmB-expressing T cells and T_RM_ cells were detected in both groups, with no significant differences in frequency between HIV-positive and HIV-negative individuals (Figure 6A). However, in line with the CXCR3-expression pattern, we observed a notable decrease in IFN-γ⁺ T_RM_ cells in both the TME and tumor of HIV-positive individuals (Figure 6B-C). These findings suggest reduced pro-inflammatory activation of tissue-resident T cell populations in HIV-associated KS, reflected by diminished IFN-γ expression.

**Figure 6.**
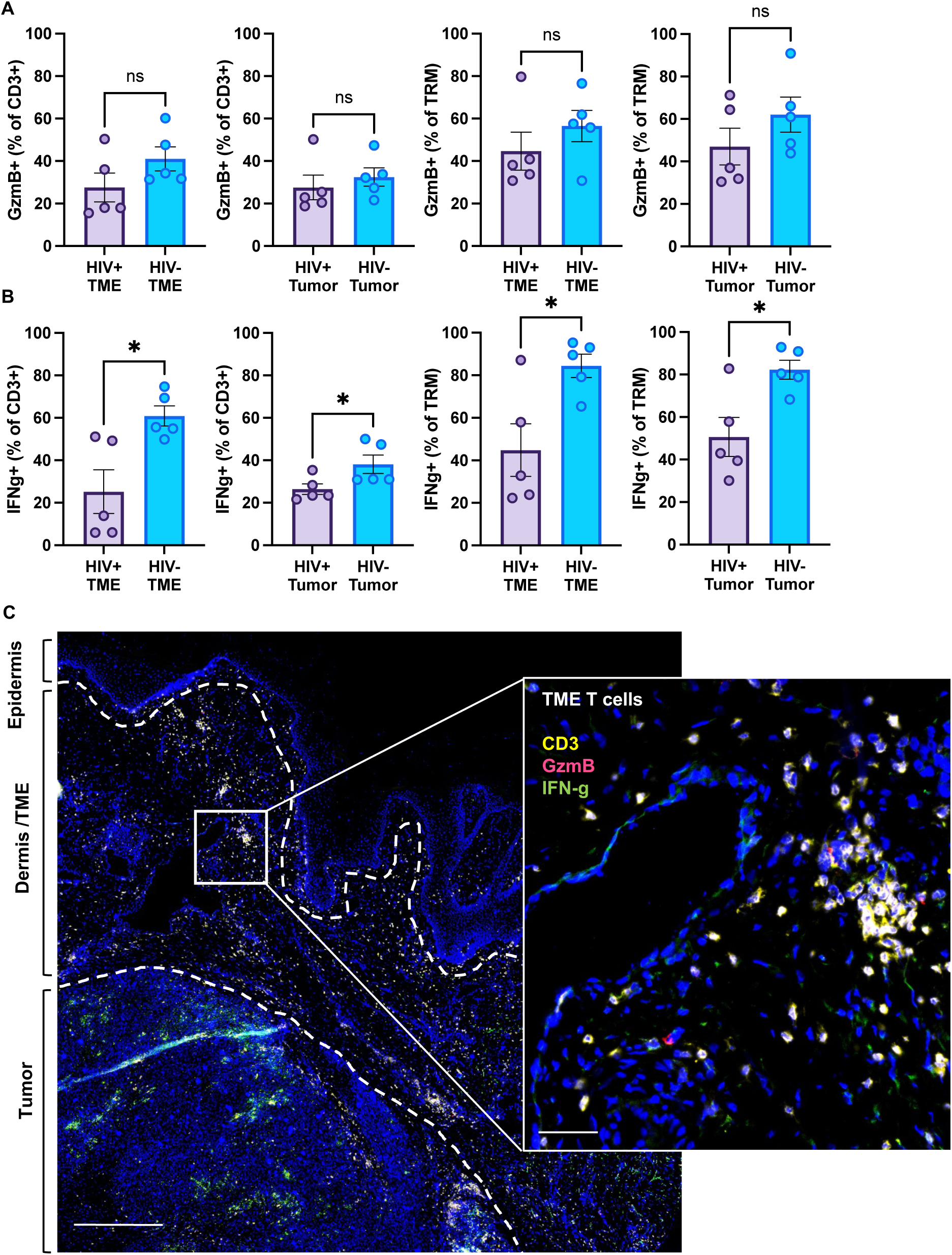
Functional polarization of T cells and TRM cells in KS tissue. (A) Quantification of GzmB-producing T cells (CD3⁺GZMB⁺) and TRM (CD3⁺CD103⁺GZMB⁺) in the TME and tumor compartments of HIV-positive (n=6) and HIV-negative (n=5) patients. (B) Quantification of IFN-γ producing T cells (CD3⁺IFN-γ⁺) and TRM (CD3⁺CD103⁺IFN-γ⁺) in the TME and tumor compartments of HIV-positive (n=6) and HIV-negative (n=5) patients. (C) Representative immunofluorescence images showing IFN-γ⁺ (green) and GzmB+ (red) CD3⁺ T cells (yellow) with nuclear counterstaining using DAPI (blue), in KS tissue sections of a HIV+ patient. Double-positive cells shown in white. Scale bar 1 (overview) = 1mm, Scale bar 2 (detail) = 100µm. In (A) and (B) data are presented as mean ± SEM. Statistical analysis was performed with unpaired Student’s t test. *P < 0.05. Data shown as percentage of T cells (CD3⁺) and TRM (CD3⁺CD103⁺).

Our analysis highlighted impairment of the local T cell type-1 activity to KS in HIV-positive individuals. To identify vantage points to increase immunogenic response in those patients, we performed putative cell-cell receptor interaction analysis from spatial transcriptomic data (Figure 7A-B). We found tumor cells to strongly interact with cells of the TME (mostly T cells), and less so with cells of the B cell lineage or structural cells (Figure 7A). This signaling involved multiple collagen subtypes, highlighting potential cell-to-cell contact (Figure 7A). Overall, the TME showed strong outgoing signaling, with a more heterogeneous pattern of pathways than those originating from tumor cells. TME-derived interactions comprised multiple extracellular matrix–receptor signaling pathways, including FN1, several collagen subtypes, and APP. These ligands predominantly engaged integrins, as well as CD44 and SDC1, on the interacting cellular compartments (Figure 7B). Interestingly, TME immune cells mainly signaled to non-tumor cell types (Figure 7B), suggesting limited direct anti-tumor activity within the TME, as well as indicating need for pharmacological promotion of interaction.

**Figure 7.**
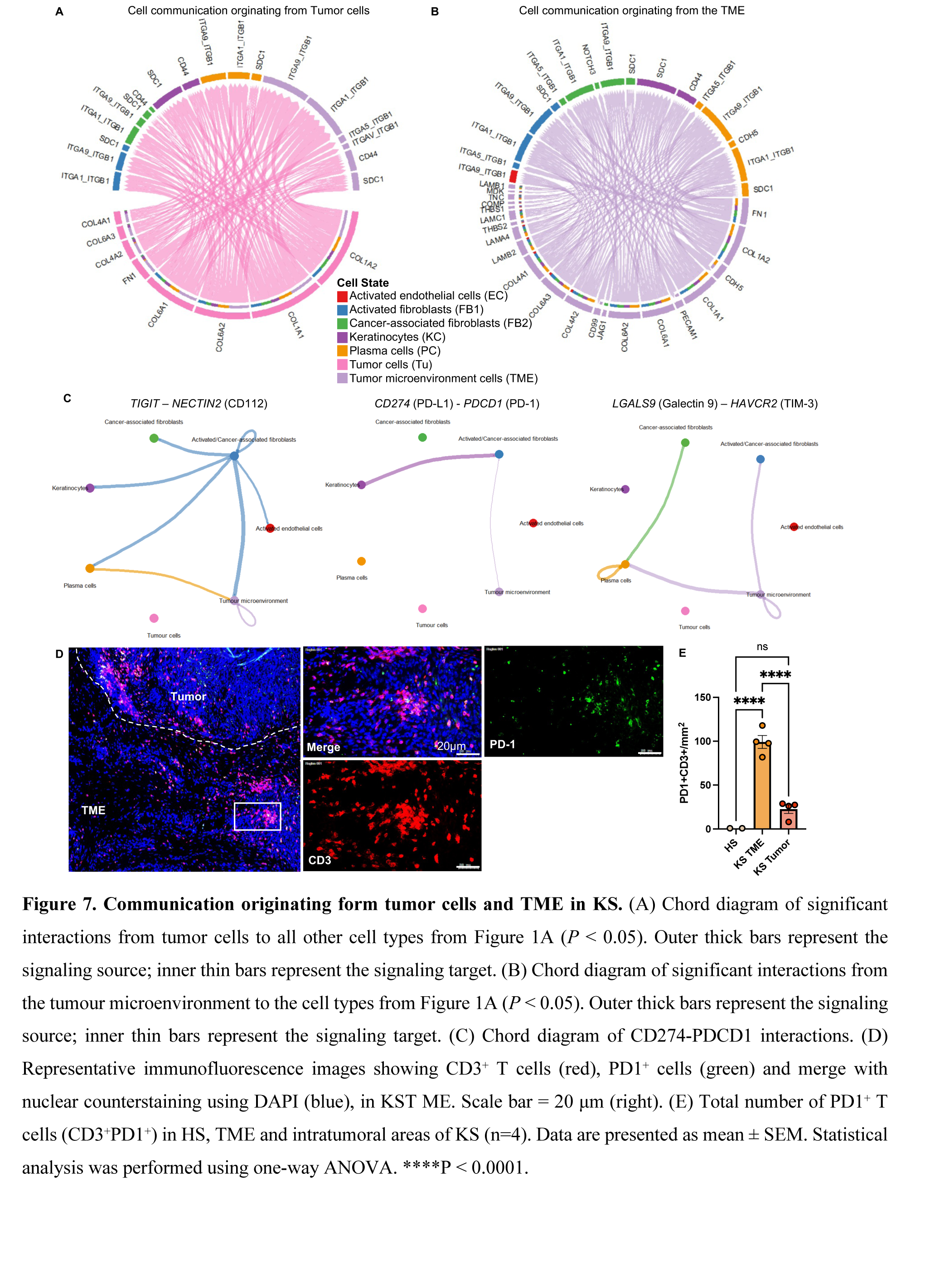
Communication originating form tumor cells and TME in KS. (A) Chord diagram of significant interactions from tumor cells to all other cell types from Figure 1A (*P* < 0.05). Outer thick bars represent the signaling source; inner thin bars represent the signaling target. (B) Chord diagram of significant interactions from the tumour microenvironment to the cell types from Figure 1A (*P* < 0.05). Outer thick bars represent the signaling source; inner thin bars represent the signaling target. (C) Chord diagram of CD274-PDCD1 interactions. (D) Representative immunofluorescence images showing CD3⁺ T cells (red), PD1⁺ cells (green) and merge with nuclear counterstaining using DAPI (blue), in KST ME. Scale bar = 20 μm (right). (E) Total number of PD1⁺ T cells (CD3⁺PD1⁺) in HS, TME and intratumoral areas of KS (n=4). Data are presented as mean ± SEM. Statistical analysis was performed using one-way ANOVA. ****P < 0.0001.

We next investigated directly analyzed interaction via major immune checkpoints described to be up-regulated in cancer (21), which revealed several putative communication hubs within peritumoral regions, including along the PD-1, TIGIT and TIM-3 signaling axes (Figure 7C, Supplementary Figure 2B). To gain more insights into potential use of immune checkpoint inhibition in KS (e.g. monoclonal antibodies against PD-1), we next investigated the protein expression of co-inhibitory receptor PD-1 on T cells which was highly increased in the KS TME compared to healthy skin (Figure 7D-E).

Overall, our data support an important role for tissue-resident, IFN-γ-associated T-cell immunity in KS and suggest that this immune program is diminished in HIV-positive individuals with low nadir CD4 counts. The observed immune landscape further provides a rationale for exploring immune checkpoint inhibition in KS.

## Discussion

The presented results demonstrate ta significant enrichment of CD8⁺ CD69⁺ and CD103⁺ T_RM_ cells in KS lesions, specifically the TME. Functionally, we demonstrated that T_RM_ cells in KS exhibit high CXCR3 and GzmB expression, indicative of a pronounced cytotoxic capacity (18). These findings are consistent with current evidence indicating that CD8⁺ T_RM_ cells play a central role in local immune surveillance of solid tumors and their presence is associated with improved clinical outcomes (22). The chemokine receptor CXCR3 is required for the migration of both tumor-specific and bystander memory T cells into solid tumors, and modulation of chemokine receptor pathways such as CXCR3 may enhance CAR-T cell therapies (23). Our data, as well as others, further implicate CXCR3 as important leukocyte recruitment factor in KS (24). The production of GzmB by T cells in the KS TME highlights their potential to directly eliminate KSHV-infected cells and is consistent with the established role of GzmB in immune surveillance of virus-associated tumors, where it is released by cytotoxic T lymphocytes and NK cells to induce apoptosis in infected and transformed cells (25).

HIV infection is known to deplete CD4⁺ T cells in tissues, profoundly impairing local immune defense (26). Delayed ART initiation leads to an irreversible depletion of CD4⁺ T_RM_ cells in the skin and mucosa, especially CXCR3⁺ T_RM_ subpopulations, despite systemic reconstitution of circulating CD4⁺ T cells. This is associated with an anti-inflammatory tissue environment that may promote the development of virus-induced malignancies, such as HPV-associated lesions (15). We found that HIV status significantly influences the extent and composition of T cell infiltration on the KS TME. As expected, we detected decreased CD4^+^ T cells in HIV-positive KS lesions, consistent with previous findings of a study investigating CD4⁺ T cells in HIV+ KS patients without highly active antiretroviral therapy (27). Furthermore, HIV-positive patients exhibited a reduced infiltration of CD8⁺ T cells in tumor tissue and in the TME. Particularly remarkable is the significantly lower number of T_RM_ cells in the TME of HIV-positive patients, with strongest effects on CXCR3⁺CD8⁺ T_RM_ cells. This is consistent with transcriptomic data showing stronger cytotoxic immune response and NK cell-mediated immunity in HIV-negative KS, whereas HIV-positive KS was characterized by alterations in growth regulation and protein kinase activity (9).

The observed increased presence of non-T_RM_ CD3⁺ cell in the TME of HIV-positive patients may reflect infiltration with bystander T cells: Chronic HIV-associated immune activation, characterized by increased T cell activation markers and pro-inflammatory cytokine expression, promotes nonspecific recruitment and accumulation of T cells in tissues, independent of their antigen specificity. Consequently, it is unlikely that a substantial fraction of CD3⁺ T cells in the TME of HIV-positive KS patients are tumor- or KSHV-specific but instead reflects systemic immune activation (28–30).

The analysis of T cell functional profiles revealed comparable frequencies of GzmB-positive T cells and T_RM_ between HIV-positive and HIV-negative cohorts, suggesting a preserved cytotoxic capacity of remaining CD8+ T_RM_ in HIV+ KS patients. A distinct difference, however, was observed in IFN-γ-producing T cells and T_RM_ cells, which were decreased in HIV-positive KS, both within the tumor and in peritumoral tissue. This consistent with reports detecting CD4⁺ and CD8⁺ T cells expressing IFN-γ in classical KS lesions, and linking robust IFN-γ and tumor necrosis factor alpha (TNF-α) secretion by KSHV-specific CD8⁺ T cells with better control of viremia and disease stability, whereas weak or less polyfunctional responses correlate with disease progression (31) (27). Overall, the presence of CD8⁺ T_RM_ cells in the KS TME of HIV-negative patients reflects intact local immunity, whereas HIV-positive patients, despite higher overall T-cell infiltration, exhibit a functionally impaired immune response with decreased CXCR3^+^CD8^+^T_RM_ and IFN-γ-producing T_RM_. Together, these differences likely contribute to the poorer outcomes in HIV-associated KS (6) (29).

Lastly, our data further confirm disrupted anti-tumor immunity in the HIV+ KS TME, which exhibited checkpoint-associated signaling pointing to a functionally impaired rather than an absent immune infiltrate. Interestingly, immune cells within the TME primarily signaled to non-tumor cell types, suggesting that these interactions may warrant pharmacological enhancement. The predominance of ECM signals suggests the presence of a stromal network that could regulate immune cell distribution and function, consistent with previous evidence that remodeling of the ECM contributes to skin tumor progression and may influence accessibility of immune cell interactions to therapeutics (32). Recent clinical data indicate that immune checkpoint inhibition with Programmed Cell Death Protein 1 (PD-1) blocking antibodies such as Pembrolizumab can induce antitumor responses in HIV-positive patients with KS (16, 33, 34). The immunomodulatory effect of PD-1 inhibition may partially reverse T cell exhaustion and promote a more effective cytotoxic and T helper-1(Th1)-polarized response, which is associated with better disease control (35). These data suggest that therapies targeting immune checkpoints should be considered a promising adjunct in the treatment of HIV-associated KS. In conclusion, our data suggest that it is not merely the presence of cytotoxic T cells that determines disease control, but rather their functional state. Particularly the quality and extent of an effective Th1-response appear to be a key determinant of immune surveillance and KS control. In contrast, chronic immune activation and functional impairment of T cells in HIV-positive patients may limit the development of a fully effective proinflammatory response and thereby contribute to a less favorable disease course.

Overall, our results suggest that an effective immune response against KS depends on both preserved cytotoxic potential of T_RM_ cells and their spatial context. From a therapeutic perspective, this provides a basis for strategies aimed at restoring effective T cell function and enhancing effective immune-tumor interactions, including checkpoint inhibition.

## Acknowledgments

The authors thank Dr. Lisa Kleissl, Dr. Thomas Krausgruber and Prof. Dr. Georg Stary for technical support with spatial transcriptomic sample submission and for providing reagents for immunofluorescence profiling. This work was in part supported by a LEO Foundation grant, received by JS (LF-OC-25-002549).

## References

1. Schneider JW, Dittmer DP. Diagnosis and Treatment of Kaposi Sarcoma. Am J Clin Dermatol. 2017;18(4):529–39.

2. Yarchoan R, Uldrick TS. HIV-Associated Cancers and Related Diseases. N Engl J Med. 2018;378(22):2145.

3. Curtiss P, Strazzulla LC, Friedman-Kien AE. An Update on Kaposi’s Sarcoma: Epidemiology, Pathogenesis and Treatment. Dermatol Ther (Heidelb). 2016;6(4):465–70.

4. Moore PS, Chang Y. Detection of herpesvirus-like DNA sequences in Kaposi’s sarcoma in patients with and those without HIV infection. N Engl J Med. 1995;332(18):1181–5.

5. Radu O, Pantanowitz L. Kaposi sarcoma. Arch Pathol Lab Med. 2013;137(2):289–94.

6. Cesarman E, Damania B, Krown SE, Martin J, Bower M, Whitby D. Kaposi sarcoma. Nat Rev Dis Primers. 2019;5(1):9.

7. Morales AE, Benson G, Glavan S, Giuliano R, Dickson MA. Fifth subtype of Kaposi sarcoma in HIV-negative MSM: a retrospective single-arm cohort study from a tertiary care center in NYC from 2000 to 2022. Oncologist. 2025;30(3).

8. Riva G, Barozzi P, Torelli G, Luppi M. Immunological and inflammatory features of Kaposi’s sarcoma and other Kaposi’s sarcoma-associated herpesvirus/human herpesvirus 8-associated neoplasias. AIDS Rev. 2010;12(1):40–51.

9. Yang J, Cali Daylan AE, Shevkoplias A, Postovalova E, Wang M, Tyshevich A, et al. Transcriptomic Profiling and Tumor Microenvironment Classification Reveal Unique and Dynamic Immune Biology in HIV-Associated Kaposi Sarcoma. Cells. 2025;14(2).

10. Lidenge SJ, Tso FY, Ngalamika O, Kolape J, Ngowi JR, Mwaiselage J, et al. Lack of CD8(+) T-cell co-localization with Kaposi’s sarcoma-associated herpesvirus infected cells in Kaposi’s sarcoma tumors. Oncotarget. 2020;11(17):1556–72.

11. Chen J, Del Valle L, Lin HY, Plaisance-Bonstaff K, Forrest JC, Post SR, et al. Expression of PD-1 and PD-Ls in Kaposi’s sarcoma and regulation by oncogenic herpesvirus lytic reactivation. Virology. 2019;536:16–9.

12. Liu W, Qin Y, Bai L, Lan K, Wang JH. Kaposi’s-sarcoma-associated-herpesvirus-activated dendritic cells promote HIV-1 trans-infection and suppress CD4(+) T cell proliferation. Virology. 2013;440(2):150–9.

13. Williams JB, Kupper TS. Resident Memory T Cells in the Tumor Microenvironment. Adv Exp Med Biol. 2020;1273:39–68.

14. Strobl J, Pandey RV, Krausgruber T, Bayer N, Kleissl L, Reininger B, et al. Long-term skin-resident memory T cells proliferate in situ and are involved in human graft-versus-host disease. Sci Transl Med. 2020;12(570).

15. Saluzzo S, Pandey RV, Gail LM, Dingelmaier-Hovorka R, Kleissl L, Shaw L, et al. Delayed antiretroviral therapy in HIV-infected individuals leads to irreversible depletion of skin-and mucosa-resident memory T cells. Immunity. 2021;54(12):2842–58 e5.

16. Lurain K, Ramaswami R, Ekwede I, Eulo V, Goyal G, Menon M, et al. Cancer Immunotherapy Trials Network 12: Pembrolizumab in HIV-Associated Kaposi Sarcoma. J Clin Oncol. 2025;43(4):432–42.

17. Krausgruber T, Redl A, Barreca D, Doberer K, Romanovskaia D, Dobnikar L, et al. Single-cell and spatial transcriptomics reveal aberrant lymphoid developmental programs driving granuloma formation. Immunity. 2023;56(2):289–306 e7.

18. Wolf FA, Angerer P, Theis FJ. SCANPY: large-scale single-cell gene expression data analysis. Genome Biol. 2018;19(1):15.

19. Jin S, Guerrero-Juarez CF, Zhang L, Chang I, Ramos R, Kuan CH, et al. Inference and analysis of cell-cell communication using CellChat. Nat Commun. 2021;12(1):1088.

20. Rauch DA, Ramos PV, Khanfar M, Harding J, Joseph A, Fahad A, et al. Single-Cell Transcriptomic Analysis of Kaposi Sarcoma. PLoS Pathog. 2025;21(4):e1012233.

21. Xiao Y, Li ZZ, Zhong NN, Cao LM, Liu B, Bu LL. Charting new frontiers: Co-inhibitory immune checkpoint proteins in therapeutics, biomarkers, and drug delivery systems in cancer care. Transl Oncol. 2023;38:101794.

22. Okla K, Farber DL, Zou W. Tissue-resident memory T cells in tumor immunity and immunotherapy. J Exp Med. 2021;218(4).

23. Sullivan PM, Reed SJ, Kalia V, Sarkar S. Solid Tumor Microenvironment Can Harbor and Support Functional Properties of Memory T Cells. Front Immunol. 2021;12:706150.

24. Uccini S, Scarpino S, Ballarini F, Soriani A, Chilosi M, Montesu MA, et al. In situ study of chemokine and chemokine-receptor expression in Kaposi sarcoma. Am J Dermatopathol. 2003;25(5):377–83.

25. Trapani JA, Sutton VR. Granzyme B: pro-apoptotic, antiviral and antitumor functions. Curr Opin Immunol. 2003;15(5):533–43.

26. Chimbetete T, Buck C, Choshi P, Selim R, Pedretti S, Divito SJ, et al. HIV-Associated Immune Dysregulation in the Skin: A Crucible for Exaggerated Inflammation and Hypersensitivity. J Invest Dermatol. 2023;143(3):362–73.

27. Guedes F, de Andrade HF, Jr., Fernandes ER, Tuon FF, Brasil RA, Pagliari C, et al. The effects of human herpesvirus 8 infection and interferon-gamma response in cutaneous lesions of Kaposi sarcoma differ among human immunodeficiency virus-infected and uninfected individuals. Br J Dermatol. 2008;159(4):839–46.

28. Nana BC, Esemu LF, Besong ME, Atchombat DHN, Ogai K, Sobgui TMP, et al. Soluble biomarkers of HIV-1-related systemic immune activation are associated with high plasma levels of growth factors implicated in the pathogenesis of Kaposi sarcoma in adults. Front Immunol. 2023;14:1216480.

29. Haas A, Zimmermann K, Oxenius A. Antigen-dependent and -independent mechanisms of T and B cell hyperactivation during chronic HIV-1 infection. J Virol. 2011;85(23):12102–13.

30. Bangs SC, McMichael AJ, Xu XN. Bystander T cell activation--implications for HIV infection and other diseases. Trends Immunol. 2006;27(11):518–24.

31. Bihl F, Berger C, Chisholm JV, 3rd, Henry LM, Bertisch B, Trojan A, et al. Cellular immune responses and disease control in acute AIDS-associated Kaposi’s sarcoma. AIDS. 2009;23(14):1918–22.

32. Fromme JE, Zigrino P. The Role of Extracellular Matrix Remodeling in Skin Tumor Progression and Therapeutic Resistance. Front Mol Biosci. 2022;9:864302.

33. Galanina N, Goodman AM, Cohen PR, Frampton GM, Kurzrock R. Successful Treatment of HIV-Associated Kaposi Sarcoma with Immune Checkpoint Blockade. Cancer Immunol Res. 2018;6(10):1129–35.

34. Delyon J, Biard L, Renaud M, Resche-Rigon M, Le Goff J, Dalle S, et al. PD-1 blockade with pembrolizumab in classic or endemic Kaposi’s sarcoma: a multicentre, single-arm, phase 2 study. Lancet Oncol. 2022;23(4):491–500.

35. Li J, Jie HB, Lei Y, Gildener-Leapman N, Trivedi S, Green T, et al. PD-1/SHP-2 inhibits Tc1/Th1 phenotypic responses and the activation of T cells in the tumor microenvironment. Cancer Res. 2015;75(3):508–18.

