## Supplementary Material for "Tissue-resident memory T cells are diminished in the Kaposi sarcoma tumor microenvironment of people living with HIV"

**Supplementary Figures and Legends**

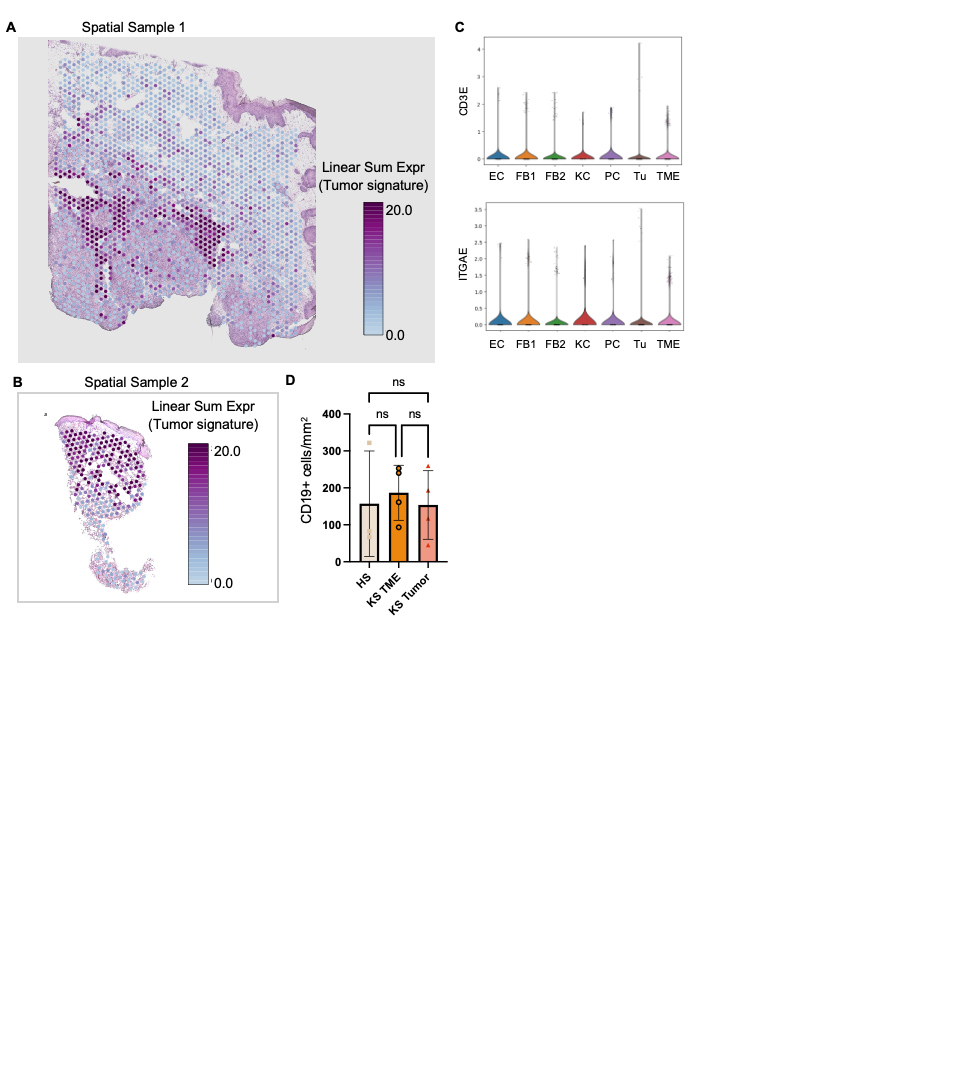

**Supplementary Figure 1:** (A-B) Expression of KS tumor signature (co-expression of LYVE1, SCN9A, CD34, PECAM1, PDPN, FLT4) in KS in Spatial sample 1 (A) and Spatial sample 2 (B). Intensity indicates average feature sum linear expression of above-indicated genes. (C) Spatial clustering (Leiden): T cells (CD3E) and T_RM_ (ITGAE) are found in the TME. (D) Quantification of CD19+ cells/mm² in healthy skin, the KS TME, and tumors. Data are presented as mean ± SEM. Statistical analysis was performed using one-way ANOVA (ns, not significant).

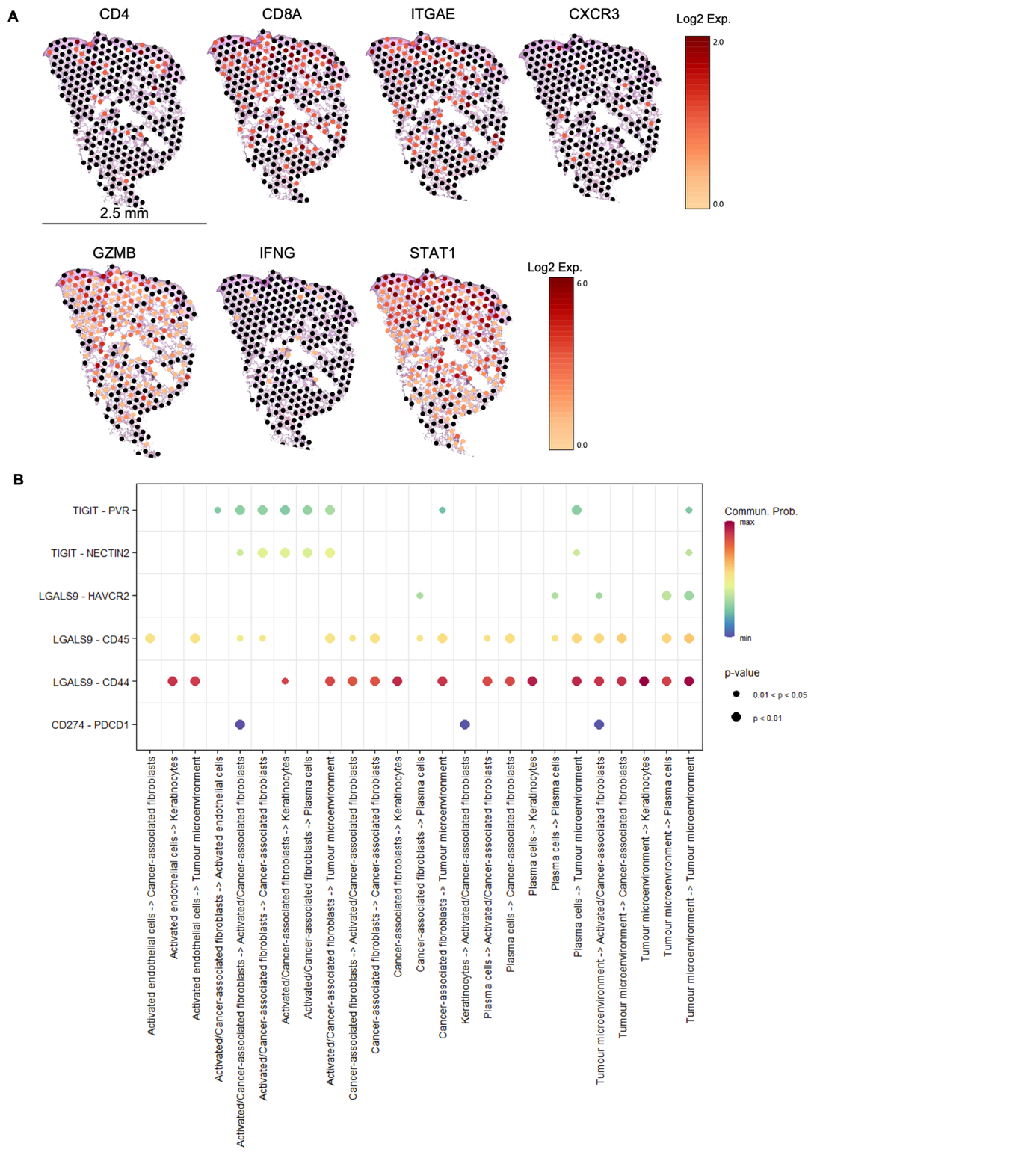

**Supplementary Figure 2.** (A) Expression of genes in KS sample 2. Intensity indicates log2 expression of each indicated gene. (B) Putative cell-cell communication of cell types identified by spatial transcriptomic profiling, focusing of inhibitory immune checkpoints.

**Supplementary Tables**

| **Pat.**  **No.** | **Age** | **Sex** | **CD4^+^/µL**  **Nadir** | **CD4^+^/µL**  **at biopsy** | **HIV c/mL at biopsy** | **HIV therapy** | **HHV-8 Status** | **KS Therapy** | **KS organ involvement** |
| --- | --- | --- | --- | --- | --- | --- | --- | --- | --- |
| P1 | 54 | m | 15 | 15 | 3 x 10^6 | BIC/FTC/TAF | positive | ART | skin |
| P2 | 51 | m | 14 | 188 | 71 | BIC/FTC/TAF | positive | ART | skin |
| P3 | 57 | w | 23 | NA | 5,623 | DTG/3TC/DRV/RTV | positive | liposomal doxorubicin | lymph nodes, skin |
| P4 | 37 | m | 129 | 129 | 9 x 10^6 | BIC/FTC/TAF | positive | liposomal doxorubicin | oral mucosa, skin, stomach |
| P5 | 55 | m | 180 | 180 | 48,978 | BIC/FTC/TAF | positive | ART | oral mucosa, skin |
| P6 | 50 | m | 49 | 221 | 129 | DOR/3TC/TDF | positive | liposomal doxorubicin | esophagus, intestine, skin |

**Supplementary Table 1.** Patient characteristics of Cohort 1, HIV-positive individuals. BIC, bictegravir; TAF, tenofovir alafenamide, FTC, emtricitabine; DTG, dolutegravir; 3TC, lamivudine; RTV, ritonavir; DRV, Darunavir; DOR, doravirine; TDF, tenofovir disoproxil.

| **Pat.**  **No.** | **Age** | **Sex** | **Comorbidities (KS etiology)** | **HHV-8 Status** | **KS Therapy** | **KS organ involvement** |
| --- | --- | --- | --- | --- | --- | --- |
| P1 | 59 | m | none (endemic region) | positive | Radiotherapy | skin |
| P2 | 73 | m | kidney transplant (tacrolimus-based immunosuppression) | positive | Radiotherapy, switch from tacrolimus to sirolimus | skin |
| P3 | 75 | w | none (endemic region) | positive | Excision | skin |
| P4 | 80 | m | none (endemic region) | positive | Excision, cryotherapy | skin |
| P5 | 55 | m | lung transplant (tacrolimus-based immunosuppression) | positive | Radiotherapy, switch from tacrolimus to everolimus, liposomal doxorubicin | lymph nodes, skin |
| P6 | 57 | m | none (endemic region) | positive | Excision, liposomal doxorubicin, radiotherapy | skin |

**Supplementary Table 2.** Patient characteristics of Cohort 2, HIV-negative individuals.
